# Do Distinct Subpopulations Signify Modes of Behavior in a Noisy Single Cell?

**DOI:** 10.1101/2025.07.22.666238

**Authors:** James Holehouse

## Abstract

Population-level distributions of fluorescence or molecule counts are often taken to reflect the behaviors of individual cells within that population. In this article, we argue that counting sub-populations can be a misleading proxy for identifying the number of behavioral modes accessible to individual cells within a system. We show that definitions of behavioral modes based on deterministic modeling can fail when fluctuations in a system’s state—or noise—become significant. In such cases, peaks in the probability distribution—emerging from stochastic descriptions—are often interpreted as substitutes for deterministically defined stable modes. However, we demonstrate that this interpretation can break down: it is possible to construct counterexamples in which two subpopulations arise from a system that supports only a single mode of behavior, driven by non-equilibrium transient dynamics. Better understanding the role of noise in transient biological randomness may allow for the discovery of novel mechanisms of regulation that are not apparent in steady state behaviors.

## I. INTRODUCTION

Populations of cells often comprise distinct subpopulations that exhibit different behaviors, characterized by variations in properties such as protein or mRNA expression, cell size, or regulatory state. These subpopulations can be seen as the modes (peaks) of a probability distribution—or histogram—over cell states, and what is seen on the population level is seen as reflective of the behaviors encoded in a single cell. When we seek to explain the occurrence of these subpopulations using mathematical modeling, we need to account for more than just the population’s differing characteristics; we also need to consider intrinsic biological randomness—or noise—present in every living system. Examining all of these factors requires that we carefully choose our models and methods of interpreting data. This introduction provides background on the two main types of mathematical models—deterministic and stochastic—and how they are used in examining the behaviors of biological systems.

### Model Types

The relationship between deterministic and stochastic modeling approaches varies depending on the particulars of the biological system under study. There are many nuances to consider, including one of particular interest to us in this paper: that deterministic modeling approaches to identifying biological behaviors break down in a wide variety of situations wherein randomness becomes important.

*Deterministic modeling* assumes that mean molecule numbers are enough to accurately specify the state of a biochemical system and that randomness is not important. Often, this is a good framework if we want to model systems without interactions in the limit of large molecule numbers or cellular volumes [1, 2]. The standard equations used in deterministic modeling are differential equations describing the time-evolution of mean molecule numbers that describe the state of the system [3, Sec. 3.1].

*Stochastic modeling* takes the role that randomness plays in biology seriously. In the context of a single cell, cellular volumes tend to be quite small, gene products (and genes themselves) often in low number, and the causes of noise can be conceptualized as coming from: (i) intrinsically random waiting times for molecular events; (ii) cell-to-cell variability [3–7]. All of these cell characteristics mean that we can best model the state of the cell as a set of random variables. This statement is verified by the multitude of methods that look at gene expression at the levels of single-cells [5, 8, 9], single molecules [10–13], and even across multiple time snapshots [14–16]. The standard equations used in stochastic modeling are coupled differential equations dictating the time-evolution of the probability distribution over all potential states of the system [1, 17].

### Seeking Explanations for Cell Behavior in Probability Distributions

Across much of cell biology, we are rarely in a situation where stochasticity can be wholly ignored, however we often use the easy interpretability of deterministic models to make claims regarding bimodality captured in biological data. For example, observations of population level protein fluorescence bimodality in [18] (see Fig. 1(a)) were predicted from a deterministic model that expressed bistability (see Fig. 1(b)). Here, deterministic bistability—the existence of two steady-state solutions of protein expression—encoded that there are two modes of behavior for this biochemical mechanism, that is, if a deterministic framework is suitable for the system under study.

**FIG. 1.**
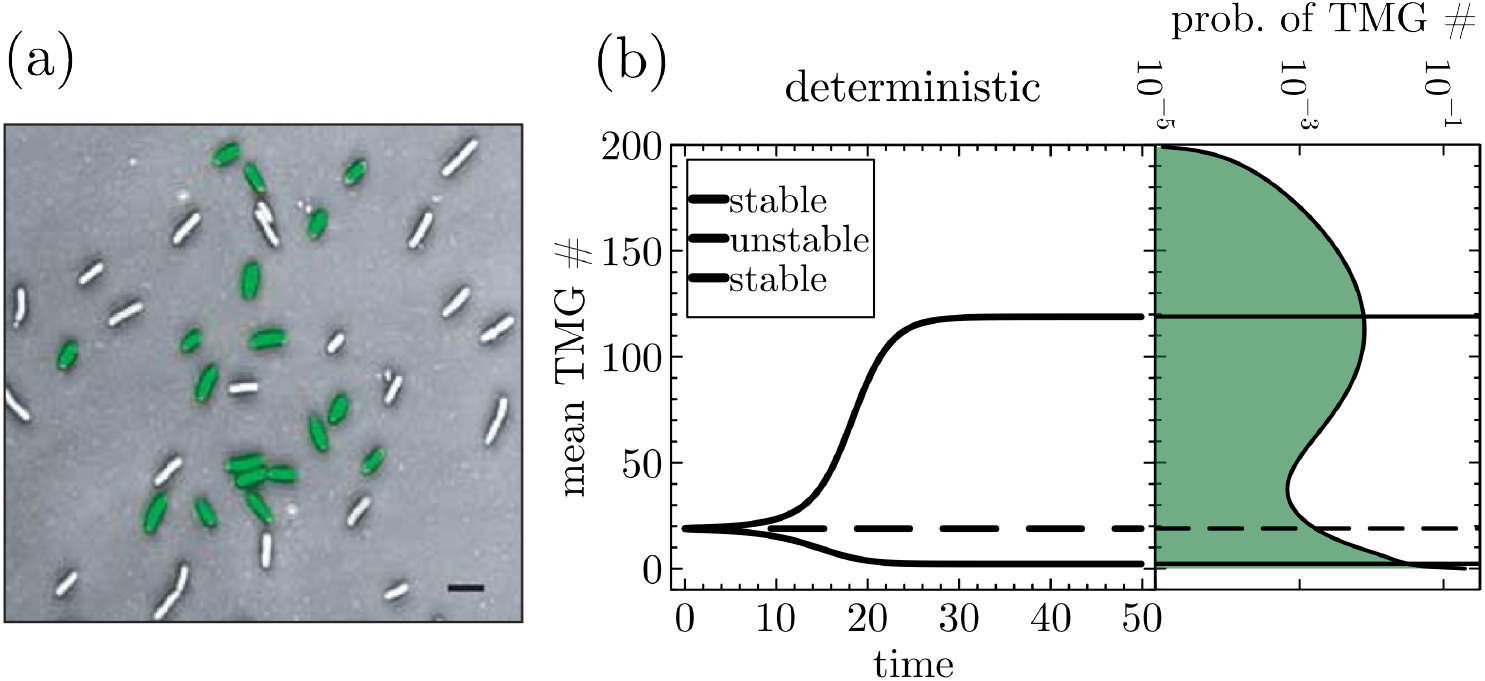
Deterministic bistability and stochastic bimodality in the *lac* system (see [18]). (a) Two distinct subpopulations of cells as observed in [18] exhibiting a bimodal distribution of fluorescence of lac expression *lac* expression (reproduced from [18, Fig. 2(a)]). (b) The left-hand plot shows bistability in the mean number of TMG molecules predicted by the deterministic model. Solid lines represent stable solutions (bistability) and the dashed line indicates an unstable solution of the deterministic equations. The right-hand plot shows the resultant bimodality in a stochastic formulation of the same model, where the vertical *x*-axis now represents TMG numbers, not mean values. For the parameters and reaction scheme used to simulate panel (b) please see SI Section S3 A.

#### Glossary

**Bistability**: When the solution of a deterministic model exhibits two unique and stable solutions for the means of the state variables (Fig. 1(b): left).

**Bimodality**: When the solution of a stochastic model is a probability distribution over the system states that has two modes/peaks (Fig. 1(b): right).

In a deterministic setting, a mode of behavior could represent the number of distinct levels of mean mRNA or protein expression, or the number of different stable arrangements of the proteome that lead to different cell types (e.g., differentiation in a Waddington landscape, see [19]). Unique sets of mean molecule numbers, for the same set of dynamical evolution equations, lead to modes of behaviors that are easily defined from a deterministic modeling perspective.

Stochastic modeling takes a different approach. Given the behavior of fluctuating amounts of mRNA and protein molecules, we are often interested in characterizing stochastic dynamics of the behavior of the cell into “modes of behavior”—broadly defined as the set of macroscopic patterns that a system can exist in. When accounting for randomness or noise becomes important, stochastic dynamics lead to a state of the cell that is continually fluctuating in time. This does not mean that regularities in cellular behavior cannot be found. For a system with stochastic dynamics, subsets of the states of a system—say a particular set of molecule numbers, rather than a deterministic mean number of molecules—define a system’s mode of behavior.

Examples of the emphasis placed on probability distribution bimodality in stochastic models are numerous [20–32] and span the literature of systems biology, synthetic biology, physics and biochemistry. In many cases, the bimodality of protein distributions involved in regulatory mechanisms can be used to explain regulatory function [22, 25], and probability distribution bimodality is seen as a functional feature of a regulatory processes in development [33, 34].

Three key insights are important to keep in mind. First, stochastic models do not bear a simple or direct relationship to their deterministic counterparts. As we show in Section III, the two kinds of models can study the same system and produce different results. Second, when these differing results occur, it is an indication that deterministic reasoning is not a reliable tool for defining behavioral modes in the system under study. Third, we should not assume that single-cell behaviors can be inferred from a population snapshot under either a deterministic or stochastic framework.

## II. PROBABILITY MODES IN SYSTEM STATES CAN BE INFORMATIVE

The practice of attributing modes of behavior to the modes of a probability distribution increased in popularity with the advent of systems and synthetic biological modeling [35–40]. In these types of modeling the identification or creation of highly heterogeneous protein expression in populations of cells was often thought to be indicated by bistability in deterministic models of mean protein expression^1^.

For the gene circuit studied in [18], the bistable predictions of a deterministic model were confirmed by a bimodal distribution of protein fluorescence states. In this case, the clear interpretation of a mode of behavior with a mode of deterministic stability—of which there are two, one of low and one of high mean protein number—was utilized to good effect. As this case shows, deterministic models can be very useful because they are easily interpretable: monostability implies a single behavioral mode, bistability implies two behavioral modes, and so on. Such identification is not clear in stochastic models since a mode of behavior is not simply identified with a mean molecule number, but a set of states.

When one utilizes stochastic as opposed to deterministic modeling approaches, it is probability modes (peaks) in the distribution over molecule numbers that are seen to provide insights into the behaviors characterizing subcellular noise. For example, stochastic models can pick up identifiable modes of behavior in systems that admit time-scale separation. The demarcation of each mode of behavior is then protected by the slow transitions between the modes, making the identification of each mode of behavior very clear, especially when one has access to time-dependent data showing the transitions between states (e.g., Fig. 2(b)). There are many examples of this type of time-scale separation in the literature: (1) reversible phenotypic switching between differing levels of transcriptional or translational control [41, 42]; (2) slow promoter kinetics [43], again leading to multimodal protein expression; (3) time-scale separation-like that approximate probability distributions of gene products through mixture distributions, such as sums of negative binomials [44, 45].

**FIG. 2.**
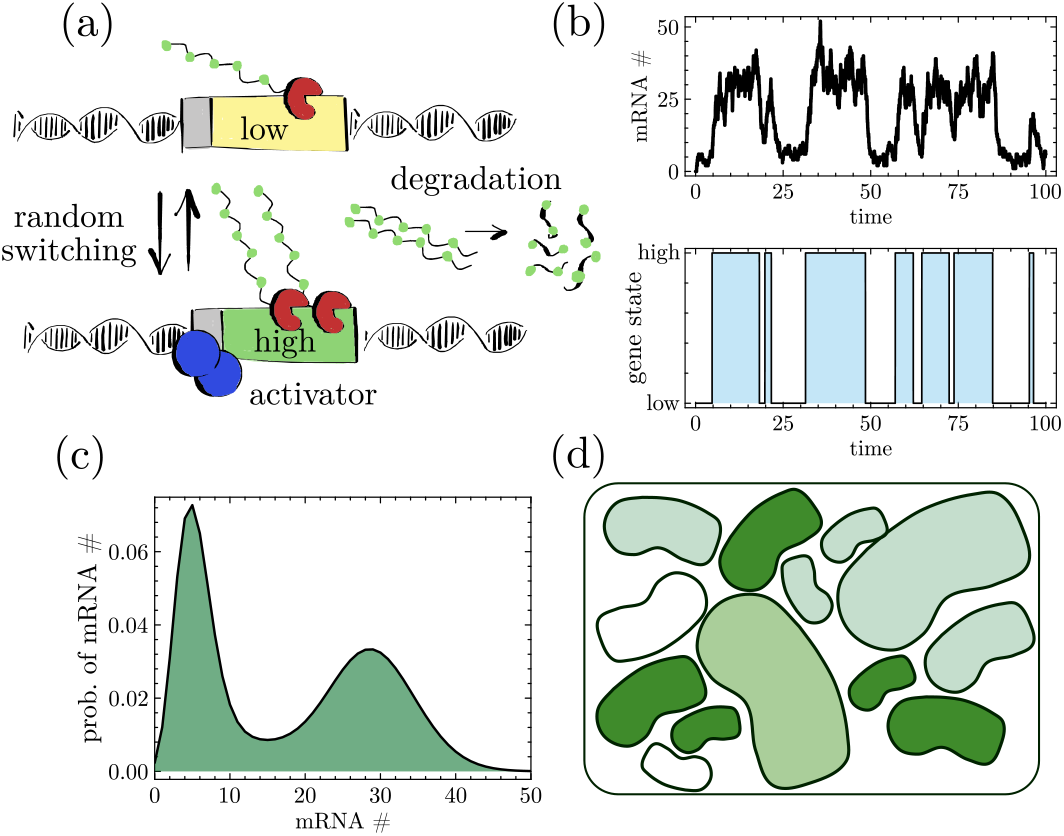
An example wherein modes of behavior correspond to modes of probability. (a) A schematic of the two-state gene system, with transcription occurring at two differing rates in each gene state (only one gene state is present at any one time), and mRNA degrades after being produced. Each mRNA is assumed to fluoresce as in a sm-FISH experiment [13]. (b) The dynamics of the mRNA number fluctuations and gene state fluctuations simulated from the stochastic simulation algorithm [46]. (c) The probability distribution over mRNA numbers. (d) A hypothetical population of cells showing two distinct populations of mRNA fluorescence, and a large variability across the population of cells. For the parameters and reaction scheme used to simulate (b) and (c) please see SI Section S3 B.

The two-state gene model (also known as the *leaky gene model* [47]) is a classic example of how stochastic modeling explains bimodal distributions arising from time-scale separation in cell populations. The two-state gene (Fig. 2(a)) can randomly switch between low and high mRNA expression states (Fig. 2(b)) upon binding to an activator or repressor [48]. In this case, modes of behavior on the level of trajectories correspond to modes of behavior in the probability distribution. This can be seen by comparing the high-low fluctuations of mRNA seen on the trajectory level (see Fig. 2(b)) with the corresponding modes of the probability distribution over mRNA numbers (see Fig. 2(c) and (d)). It is this type of mechanistic explanation of bimodality—linking probability modes to trajectory-level behaviors—that often dominates the modeler’s mindset: where there is a mode of probability there is a mode of behavior. Often, this can be a useful heuristic. But it can also be misleading, as we show in the next two sections, where we consider systems for which deterministic arguments can no longer be relied upon, and where transient effects can lead to bimodal distributions of molecule numbers without two behavioral modes.

## III. KEIZER’S PARADOX: WHEN DETERMINISTIC DEFINITIONS OF BEHAVIORAL MODES FAIL

As shown in the previous section, modes of behavior in the deterministic sense are easy to count—they simply correspond to the number of stable values of mean mRNA or protein number. In an ideal setting, deterministic and stochastic modeling approaches lead to convergence on the modes of behavior in a given system. For example, if one has a deterministic model exhibiting bistability in the steady-state regime, then one might expect that the equivalent stochastic model would have a probability distribution with two modes, each mode being centered over a mean value predicted from the deterministic model. Although this matching does frequently occur (e.g., Fig. 1(b)), in the general case they do not match. For example, in Fig. S1 we show that the system studied in Fig. 1 can exhibit bistability without bimodality, for a different set of parameters. This kind of result is more likely to occur when molecule numbers are low and when steady-state conditions cease to hold. Under such conditions, the richness of stochastic dynamics leads to fluctuations that are non-Gaussian. In addition, standard approximations break down, such as Taylor expanding about deterministic mean values (i.e., the linear-noise approximation) [1].

The phenomenon of deterministic and stochastic model mismatch is known as Keizer’s Paradox—a term conceived in [49], based on [50]. For Keizer, this finding was in the context of the reaction scheme *A* + *X →* 2*X*, 2*X → E*. This scheme has an absorbing state of zero *X* molecules under a stochastic formulation, whereas the deterministic approach gives a single stable non-zero mean value of *X* [50, p. 164-165]. This is the paradox: *that deterministic and stochastic formulations of the same process can give wildly different qualitative results*. Other examples of Keizer’s paradox include the previously discussed two-state gene, which is always deterministically monostable even in regimes that have stochastic bimodality, genetic autoregulation, and the Moran model describing genetic drift. The Moran model is particularly interesting—the deterministic prediction becomes the least likely scenario in the stochastic case—i.e., deterministic and stochastic models give exactly the opposite result [51–54]!

The bistability–bimodality relationship breakdown carries an important takeaway: we should be cautious about whether finite-molecule effects or small-volume noise might substantially impact the observed phenomenology. If this is suspected, then we should abandon deterministic models and utilize a stochastic approach that is designed to deal with variability and finite molecule number effects^2^. In the absence of deterministic models that provide easily interpretable modes of behavior, counting the number of probability modes in stochastic models is the go-to heuristic—an intuitive but not always reliable assumption that we examine in the next section.

## IV. BIMODALITY WITH TRANSIENT—NOT BEHAVIORAL—MECHANISM

The two-state gene studied in Fig. 2 illustrates a case where two distinct behaviors at the trajectory level are reflected in the distribution over molecule numbers. This follows the standard heuristic, wherein probability modes—from the population level—correspond to behavioral modes as seen on a single-cell trajectory level. In this section, we present an example where this relationship between trajectory-level behavior and distributional modes breaks down.

**Transient Bimodality in Cell Biology**

When the number or fluorescence distribution over a particular mRNA or protein in a population of cells expresses two modes while relaxing towards an asymptotic state. *Transient bimodality is not due to competing mechanisms of steady state behavior*.

Consider an enzyme, *E*, that forms a 3*E* oligomer which processes a substrate, *S*, into product, *P*, via the Michaelis-Menten-like scheme *S* + 3*E* ⇌ *C → P* + 3*E*. Note that changes in the number of subunits making up the oligomer do not affect the phenomenology of the results that follow, and the 3*E* oligomer was simply chosen as a plausible enzymatic mechanism [55, 56]. One can see this as a catalytic process of converting *S* → *P*; starting with *M* substrate molecules the process will end with *M* product molecules. There is a single mechanism of transforming *S* → *P*. Correspondingly, the substrate number trajectories from the initial to the final state seem to be described by a single trajectory, with some noise about the mean trajectory (see Fig. 3(d)). Following the standard heuristic, one might predict that this process has a single mode of probability, starting from the state of *M* substrate molecules, and ending with a single mode of *M* product molecules. However, this is not what is observed in the general case.

**FIG. 3.**
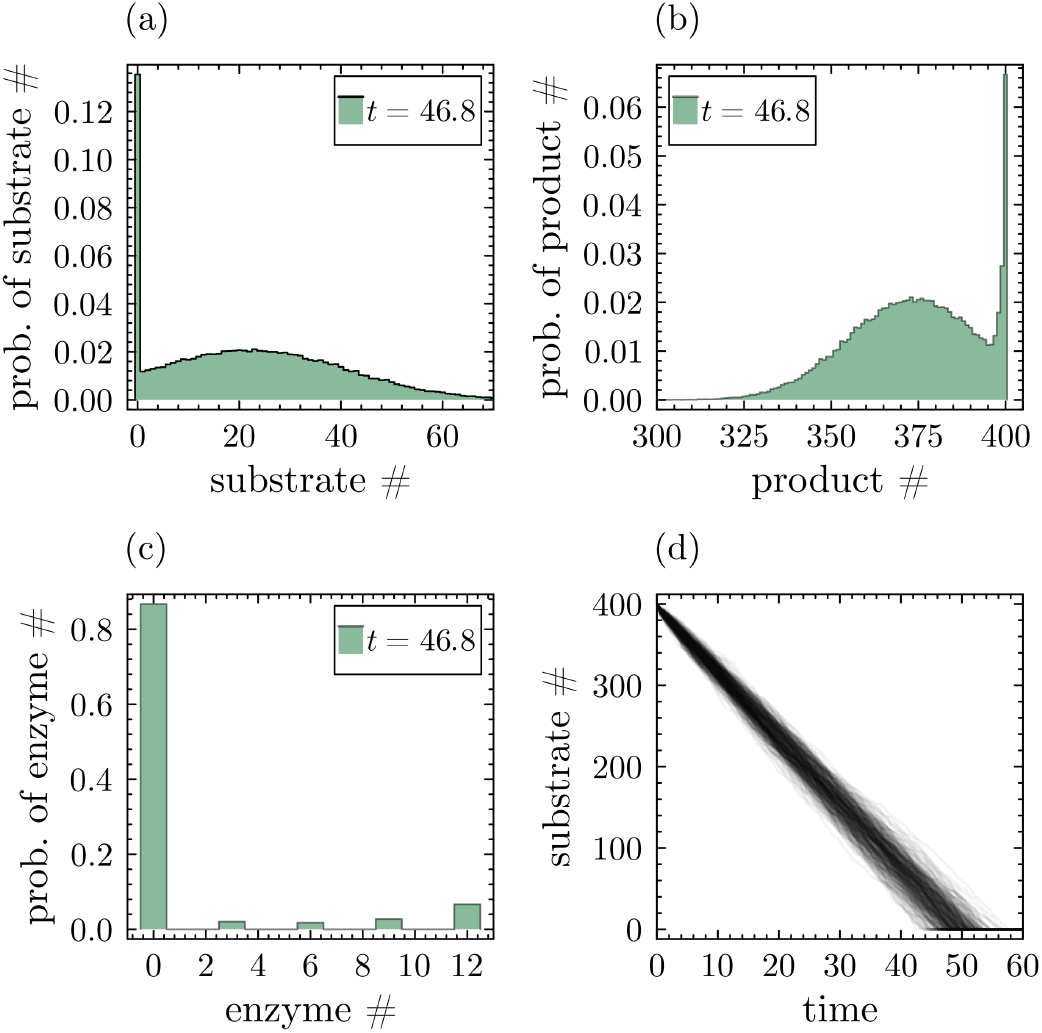
Transient bimodality in enzyme kinetics; *S* + 3*E* ⇌ *C* → *P* + 3*E*. (a) At intermediate times the distribution over substrate becomes highly bimodal even though there is only a single mechanism of substrate-to-product decay. In (b) and (c) we show that the distributions over product number and enzyme number are also bimodal. (d) Shows the decay of the substrate number in time, and importantly only a single mode of behavior with small intrinsic fluctuations. See Section S2 for the parameters that were used in these simulations.

In Fig. 3, we show an example of this enzymatic process giving rise to *transiently bimodal* distributions in the substrate (Fig. 3(a)), product (Fig. 3(b)) and enzyme (Fig. 3(c)) molecule numbers. That is, a single biological behavior is expressed through a population level probability distribution with two modes. It is this contrast between what is seen on the level of trajectories, and what is seen in the distribution over system states that should be counter-intuitive to the standard heuristic of assigning probability modes to behaviors. This is a significant conceptual issue, *since a direct observation of two subpopulations of cells here refers only to the transient intrinsic noise between cells, and is not related to there being two modes of behavior encoded in the dynamics of a single cell*. One can see this as a breakdown between the Eulerian, fixed coordinate, perspective— wherein two modes of behavior are observed—and the Lagrangian, trajectory-based, perspective—wherein only a single mode of behavior is observed.

Transient bimodality has become recently highlighted in the biological literature, having been predicted in systems as wide-ranging as standard Michaelis-Menten enzyme kinetics [57, 58] and genetic auto-regulation [59– 61]. Identification of transiently bimodal phenomena goes back to physics literature showing transient bimodality in the dynamics of lasers [62] and the theory of “chemical explosions” [63, 64]. In their classic text [65, p. 174-179], Nicolis and Prigogine discuss the origins of such explosive transient behavior for general exothermic reactions. Therein, they show how transient bimodality can be a very long-lived phenomenon [65, Fig. 79] which they describe as a “bifurcation behavior in time”. This is important, since often in cell biology we utilize the idea of the “biological steady state” [66], but in the presence of long-lived transient behaviors one cannot be sure that the heterogeneity and multi-modality seen in a population of cells is truly the result of competing behavioral modes. Several studies have attempted to explain the conditions necessary for transient bimodality—that it is the result of an initial condition interacting with a steady state or an absorbing state, in a situation where there is a “long induction stage followed by a fast switching to its steady state” [65, 67]. However, this explanation of transient bimodality does not explain the case of enzyme kinetics, which expresses transient bimodality, and does not exhibit fast switching to its absorbing state [57].

Note that transient bimodality is quite a robust phenomenon, and in Fig. S2 we show that even quite substantial noise^3^ in the kinetic parameters of the enzyme kinetic scheme explored in Fig. 3 still results in transient bimodality in substrate, product and enzyme numbers (in the case of enzymes, the transient bimodality is amplified by cell-to-cell variability). More work needs to be done to confirm the robustness of transient bimodality in the general case. There are also examples of bimodality arising purely from cell-to-cell differences (known as extrinsic noise [6]) which is unrelated to bimodality arising from competing behavioral modes intrinsic to a molecular process [68, 70–73].

## V. DISCUSSION

In this article we have highlighted the disconnect between deterministic and stochastic modeling approaches for the dynamics of processes inside a single cell, and further shown that the modes of probability distributions at the population level do not necessarily correspond to behavioral modes in the dynamics of a single cell. This can easily result from long-lived transient behaviors, which has long been recognized as important in the econophysics literature [74, 75]. If we take the idea of cells not being in a steady state seriously, such time-dependent effects should also be considered in mathematical models of the cell, and in biology more widely.

The robustness and origin of the phenomenon of transient bimodality has yet to be explored widely in the literature, and in general, such effects are often assumed to be unimportant. This article makes the case that these transient effects are very important, as the misidentification of the number of behavioral modes constitutes a significant qualitative error in the mathematical modeling of biological systems. Transient bimodality has yet to be clearly identified experimentally in biological systems, but it has been observed in optical devices [76, 77], impressive due to the long metastable times that transient bimodality can exhibit.

This article will perhaps allow us to see existing biological data in a new light: that it is on the level of multiple time-point trajectories in single cells, and not on the level of the distribution over cell states in a population, that is the clear indicator of a behavioral mode. Other recent studies have highlighted the utility of a trajectory-based approach to distinguishing between two processes that is not possible using binned histograms [78]. Our results will hopefully inspire experimentalists to search for transient bimodal behaviors in novel settings, and to hypothesize on the role it could play in biological mechanisms. In particular, the phenomenon of transient bimodality gives a purely non-equilibrium relaxation-based approach to inducing distinct cellular subpopulations that does not require resource-expensive regulatory architectures.

When bimodality is observed in a stochastic model, a question arises in how that bimodality might be biologically useful. Perhaps the question should be: can we see the origins of bimodality in the trajectories of the cell’s state variables, and if not, how can we explain its occurrence?

## ACKNOWLEDGMENTS

This work was supported by a Lou Schuyler grant from the Santa Fe Institute and National Science Foundation grants DMR-191073 and 2133863. JH thanks Ramon Grima for many insightful conversations, Kaan Öcal and Augustinas Sukys for comments, and Amy Clipp for writing advice.

## Supplementary Information

### S1 DEFINITIONS

For the reader’s ease, we provide a glossary of terms used in the main text that may be unfamiliar to those from less mathematical backgrounds.

- **State variable**: a property of a system, e.g., the number of molecules of a given protein, that tells one about the biochemical state of part of the system of interest.
- **System’s state**: the set of all state variables necessary to characterize the system of interest.
- **Mode of behavior** (behavioral mode, metastable state): a set of states—e.g., set of protein numbers—that are frequently visited, wherein the probability of transitioning between states in the mode of behavior is far greater than the probability of transitioning outside of it.
- **Mode of a probability distribution** (probability mode): a peak or maxima in the probability distribution over a set of biological states.
- **Deterministic model**: a set of equations modeling the mean of the state variables assuming no fluctuations [3, Sec. 3.1].
- **Stochastic model**: a set of equations that model the transitions between all possible arrangements of the system’s state variables [3, Sec. 3.2].
- **Bistability**: when the solution of a deterministic model exhibits two unique solutions for the means of the state variables.
- **Bimodality**: when the solution of a stochastic model is a probability distribution over the system states that has two modes/peaks.
- **Multi-stability**: when the solution of a deterministic model exhibits more than one stable and unique solutions for the means of the state variables.
- **Multi-modality**: when the solution of a stochastic model is a probability distribution over the system states that has more than one mode/peak.
- **Steady-state dynamics**: when the dynamics between the states of the system occur in the regime in which time-averages over the states of the system are constant.
- **Transient dynamics**: when the dynamics between the states of the system occur in the regime in which time-averages over the states of the system are not constant.

### S2 THE STATISTICAL MECHANICS OF MODES OF BEHAVIOR

The standard practice of attributing *modes of probability* to *modes of behavior* arises from non-equilibrium statistical mechanics, in particular the Fokker-Planck equation [1, 17, 79]. The Fokker-Planck equation provides a general stochastic modeling framework that can be used to describe the properties of chemical reaction networks inside the cell, and it assumes that molecule numbers are continuous. The solution of the Fokker-Planck equation is the probability distribution over the states of the system at a given time (often in the steady-state regime), and the peaks of this distribution can be related to the minima of a potential that associates an energy-like value to a given configuration over states [17, p. 133] [79, Sec. 6.3-6.5]. If there are two minima of the potential describing this probability distribution, then one can show that the time to transition between the two states defined by these minima is exponential in the potential difference between the maximum and the minimum of the potential [17] [79, p. 168-169]. Hence, the transition time between the two states defined by the minima is exponentially long, and these two minima can be said to represent modes of behavior—a set of states of the systems in which transitions occur much faster inside of the mode of behavior than across states located in different behavioral modes.

For a summary of the framework underlying steady-state modes of behavior in systems without discrete molecule number dynamics (akin to chemical reactions in large volumes) please see [80]. When the Fokker-Planck equation can be solved in time, and not simply in the steady state [79, Eq. (6.129)], it is possible to see the time-dependence of the potential function in time. However, such exact solutions of the Fokker-Planck equation are generally quite rare outside of one-dimensional (single molecular species) cases.

### S3 SUPPLEMENTARY INFORMATION FOR THE FIGURES

#### A. Figure 1

The deterministic rate equations used to model the *lac* system in [18] are given by,

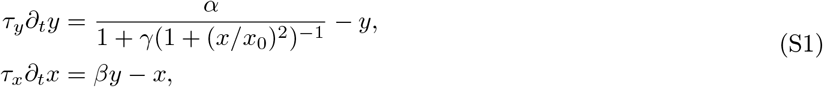

in which *x* and *y* are the respective concentrations of intracellular TMG and LacY, and all other parameters are kinetic parameters whose mechanistic interpretation is given in [18, SI “Positive feedback and bistability”]. Although the focus of [18] was in the fluorescence of LacY, here we looked at the number of TMG molecules from this reaction scheme. Note that when *x* expresses bistability, *y* does too. The corresponding stochastic reaction scheme is then given by,

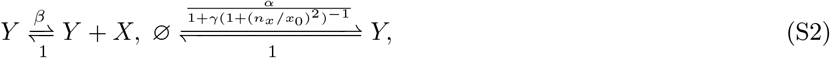

when *τ*_*x*_ = *τ*_*y*_ = 1 and in which *n*_*x*_ is the number of molecules of *X* and where we assume the volume of the system to be 1. Note that this stochastic formulation of the deterministic rate equations is a common heuristic used in the literature, and its validity is generally well-approximated in the regime when molecule numbers are large. For more details please see [81]. In both Fig. 1(b) and Fig. S1 we used the finite state projection algorithm to determine the probability distributions over TMG number [82]. The deterministic rate equations were evaluated using Mathematica’s function NDSolve.

For Fig. 1(b) the parameters that we use are: *α* = 20, *β* = 7, *γ* = 70, *x*_0_ = 6. For Fig. S1 the parameters that we use are: *α* = 10, *β* = 6, *γ* = 20, *x*_0_ = 6. Supplementary Figure Fig. S1 shows a clear case in which the deterministic model expresses bistability and the stochastic model expresses only mono-modality.

**FIG. S1.**
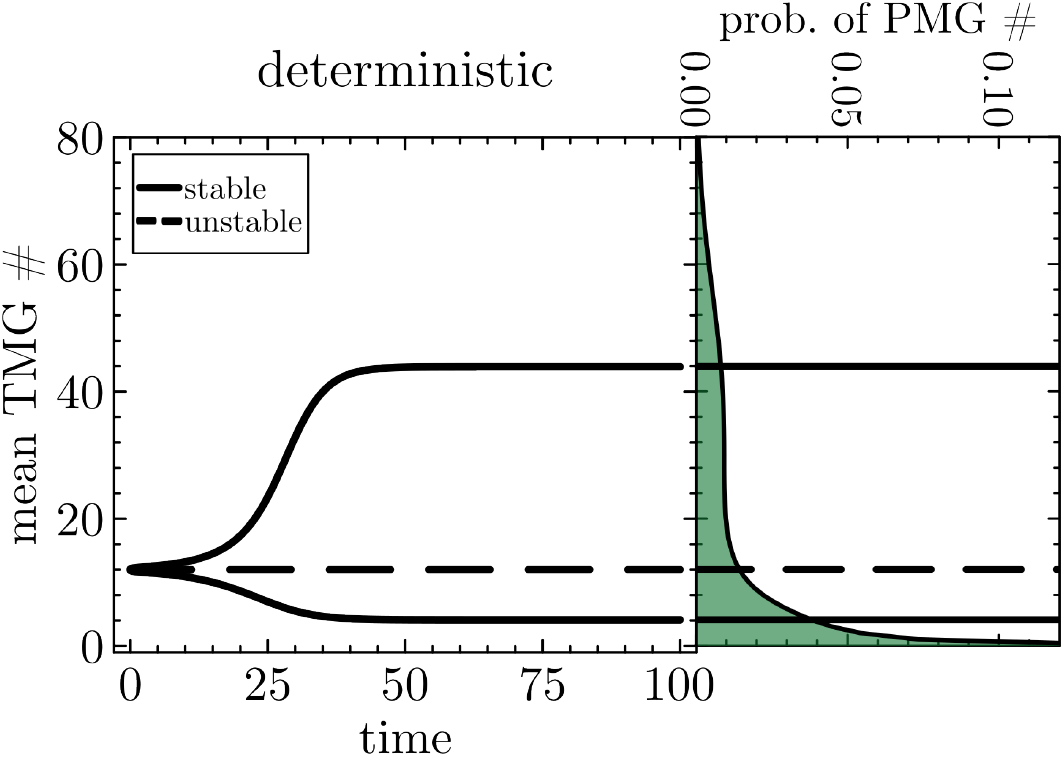
Example wherein the reaction scheme in Eq. (S2) expresses bistability without bimodality.

#### B. Figure 2

The classic two-state gene model of transcription is given by the following set of reactions,

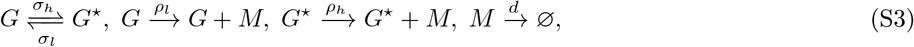

in which *G* is the low expression gene state, *G*^*⋆*^ is the high expression gene state, *M* is the mRNA and *M →* ∅ represents the degradation/dilution of mRNA. We simulated this reaction scheme using the stochastic simulation algorithm (in Julia [83]) to produce Fig. 2(b) using parameters: *σ*_*l*_ = 0.1, *σ*_*h*_ = 0.1, *ρ*_*l*_ = 5, *ρ*_*h*_ = 30 and *d* = 1 [46]. We used the same set of parameters in the finite state projection to produce Fig. 2(c) [82]. B Figure 2 3

#### C. Figure 3

The reactions defining the enzyme kinetic scheme in Figure 3 are given by,

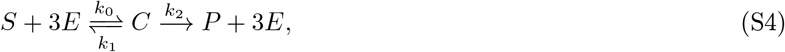

in which *k*_0_, *k*_1_ and *k*_2_ are the respective kinetic rates for enzyme-substrate binding, enzyme-substrate unbinding, and the rate of converting the enzyme-substrate complex successfully into product molecules. The distributions in Fig. 3 are the result of binning over 10^5^ individual trajectories at time *t* = 46.8 (chosen to emphasize the transient bimodality across the three species with a linear *y*-axis). Each trajectory started with 400 substrate molecules and was simulated using the stochastic simulation algorithm [46]. In Fig. 3 the chosen parameters are: *k*_0_ = 100, *k*_1_ = 1 and *k*_2_ = 2, and we simulated the process assuming there are initially 12 free enzyme molecules.

**FIG. S2.**
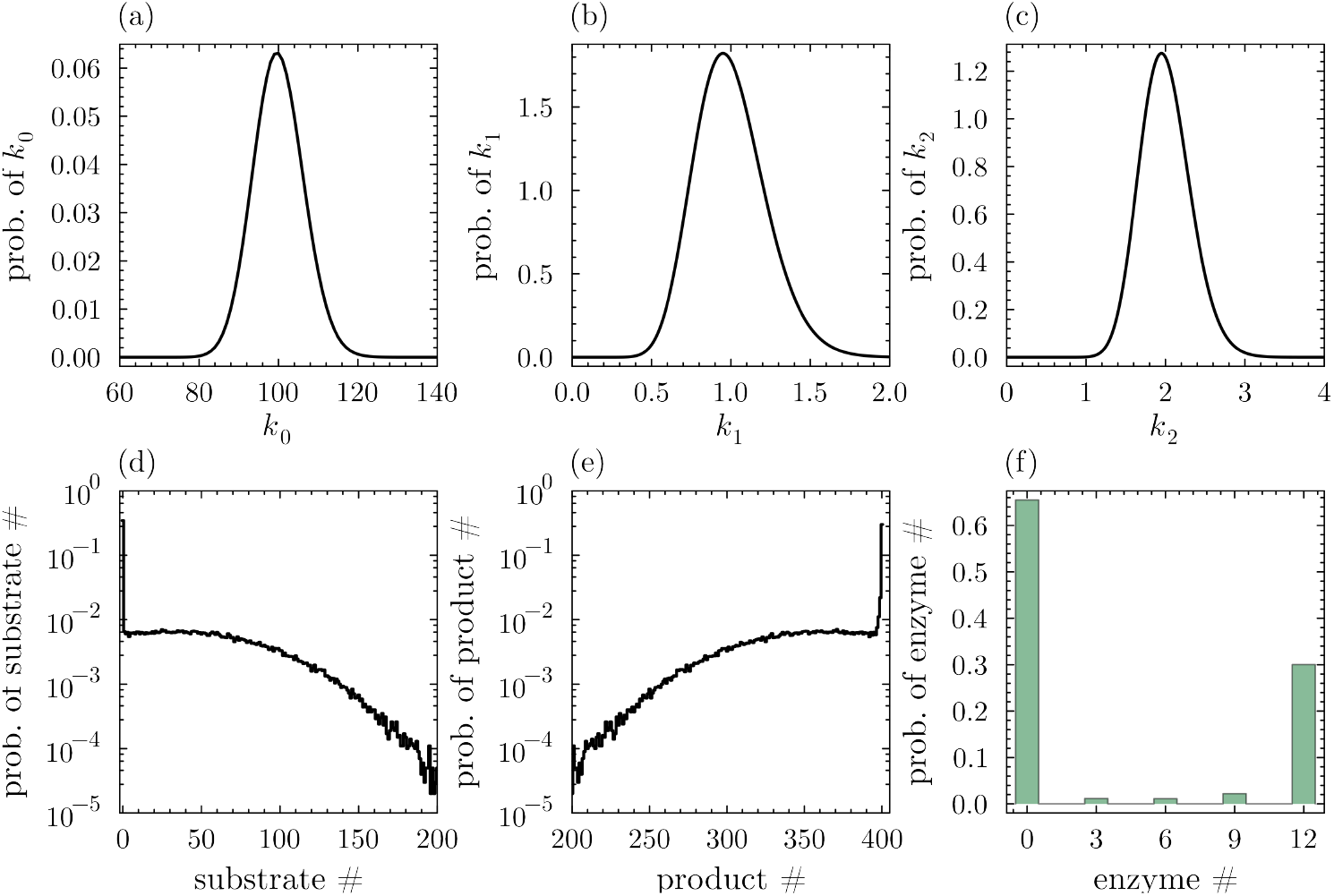
Figure showing the robustness of transient bimodality to parameter noise. (a)-(c) Show the distributions from which *k*_0_, *k*_1_ and *k*_2_ were drawn for each individual trajectory (all gamma distributions). (d)-(f) Show the resultant probability distributions over molecule numbers at *t* = 46.8 (same time as in Fig. 3(a)-(c)).

In Fig. S2, for the same parameters as in Fig. 3, we show the robustness with respect to noise in the kinetic rate parameters across different trajectories. For each trajectory, the distributions in Fig. 3(a)-(c) were sampled to give values of *k*_0_, *k*_1_ and *k*_2_; in this way we aimed to simulate the effect of cell-to-cell variability along conditions such as cell volume, pH and concentrations of enzymatic co-factors. Fig. 3(d)-(f) show that even with this parameter noise transient bimodality is still found, and with a much longer tail of substrate and product distributions. This shows that transient bimodality is a relatively robust phenomenon, at least in this case, although further work needs to be done to show the robustness of transient bimodality in the general case.

## Footnotes

1 For the statistical mechanical origin of the identification of modes of behavior with modes of probability, see Section S2.

2 Note, this is the opposite of Keizer’s view that the stochastic formulation of molecular kinetics can lead to results that are “physically meaningless” when compared to the solution of the deterministic formulation (see [49]).

3 Such extrinsic noise can arise from cell-to-cell differences in pH, cell volume and enzymatic co-factors [68, 69].

